## SupplementaryMaterial for "Distinct cytosolic complexes containing the type III secretion system ATPase resolved by 3D single-molecule tracking in live *Yersinia enterocolitica*"

**Figures – Supplemental**

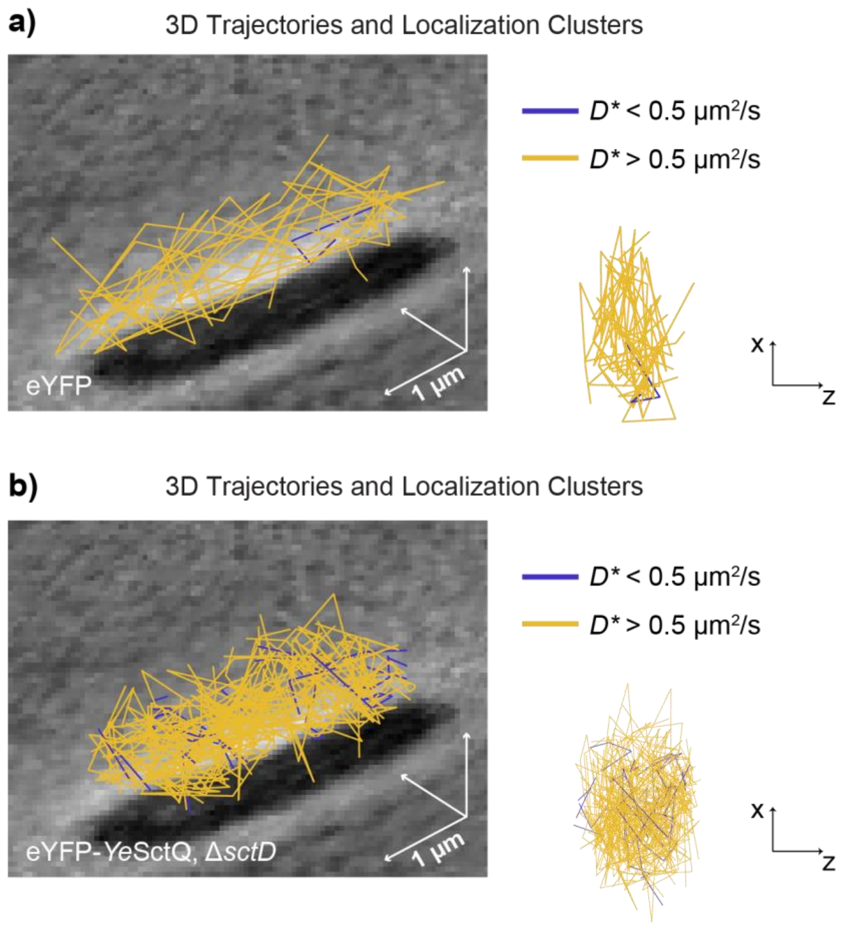


**Supplementary Figure 1:** a) 3D trajectories for eYFP under secretion active conditions. b) Trajectories classified as with $D^* < 0.5$ µm$^2$/s are not localized to the cell membrane. b) 3D trajectories for *Ye*SctQ Δ*sctD* under secretion active conditions. Trajectories classified as with $D^* < 0.5$ µm$^2$/s are not localized to the cell membrane.

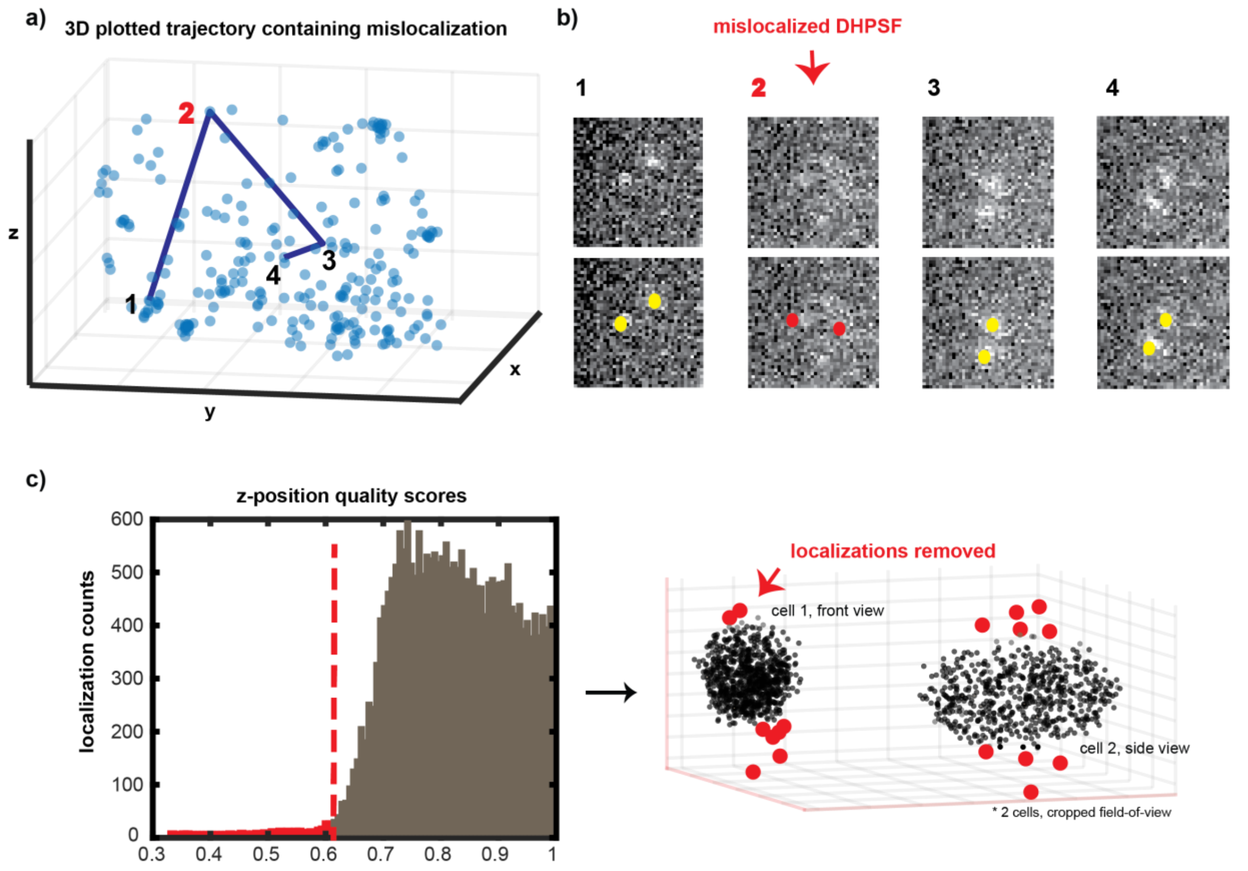


**Supplementary Figure 2:** a) 3D scatterplot of *Ye*SctQ localizations in a single bacterial cell. Localizations of a putative single-molecule trajectory are connected by blue lines. However, the second localization corresponds to an incorrect localization. b) Individual camera frames giving rise to the connected localizations shown in panel a. The second frame shows only a dim fluorescence signal due to fluorophore blinking, which leads to mislocalization of the emitter. c) z-position quality scores of all localizations detected in $N \sim 40$ cells across the field-of-view. *z*-position quality score of the *i*th emitter is computed as $QS_i = exp(-a * |z_i - \langle z \rangle|)$, where $a = 0.0008$ and $\langle z \rangle$ is the average z-position of all emitters.

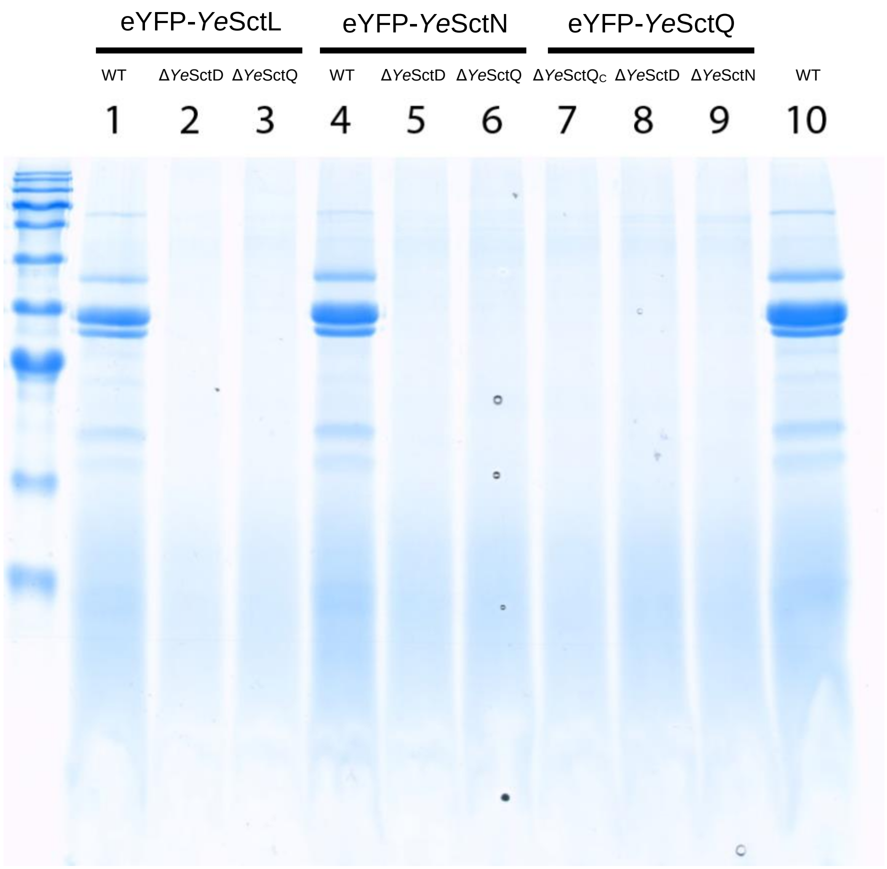


**Supplementary Figure 3:** Secretion profile for mutant strains expressing eYFP-labeled cytosolic injectisome proteins in place of the unlabeled protein. eYFP-*Ye*SctL and eYFP-*Ye*SctN both show near WT levels of secretion in the WT background. The deletion of any one (cytosolic) injectisome protein abolishes secretion, as observed previously[36]. Also of note is the loss of secretion for eYFP-*Ye*SctQ in the Δ*Ye*SctQ$_C$ background, further confirming the functional relevance for *Ye*SctQ$_C$ for secretion.

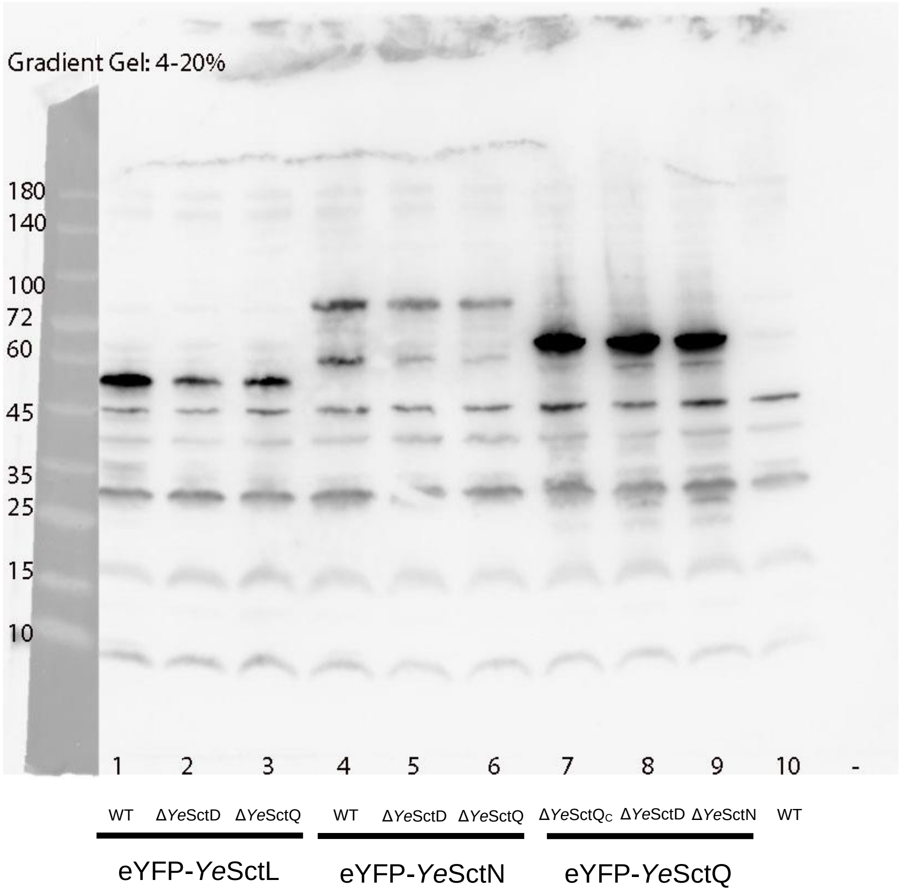


**Supplementary Figure 4:** Expression levels and stability of the used fusion proteins. All fusions proteins seem to be expressed to a similar degree (YscQ > YscL > YscN, as expected) at the expected MW. No visible degradation is evident for eYFP-YeSctL, some degradation for eYFP-YeSctN (~ 60 kDa, ~20-40% of intensity of full-length band, and very weak degradation for eYFP-YeSctQ (weak band at around 55-60 kDa).

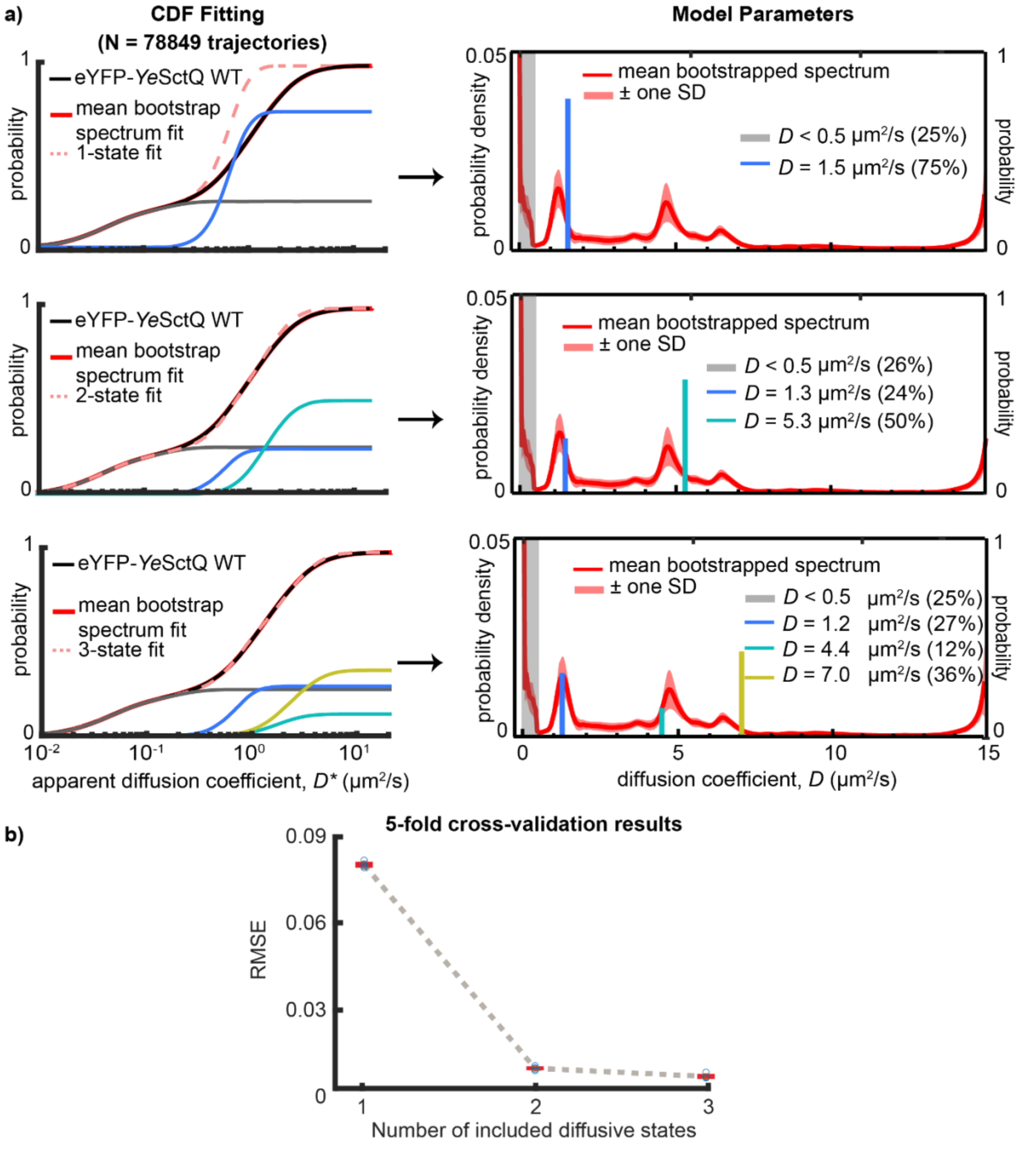


**Supplementary Figure 5:** a) Fitting model comparison for eYFP-*Ye*SctQ single-molecule tracking data. Individual rows correspond to 1-, 2-, and 3-state models, respectively. A 2-state model converged on $D \sim 1.3$ and 5.3 μm$^2$/s, which agrees well with peaks in the diffusion coefficient spectrum. b) The 2-state model is also supported by 5-fold cross-validation analysis, which shows that a 2-state model does not overfit the data. A 3-state model could also have been chosen based on the cross-validation analysis alone. However, the 3-state model includes diffusive states at $D \sim 7.0$ μm$^2$/s with an overly large population fraction of 36%, which does not match the diffusion coefficient spectrum.

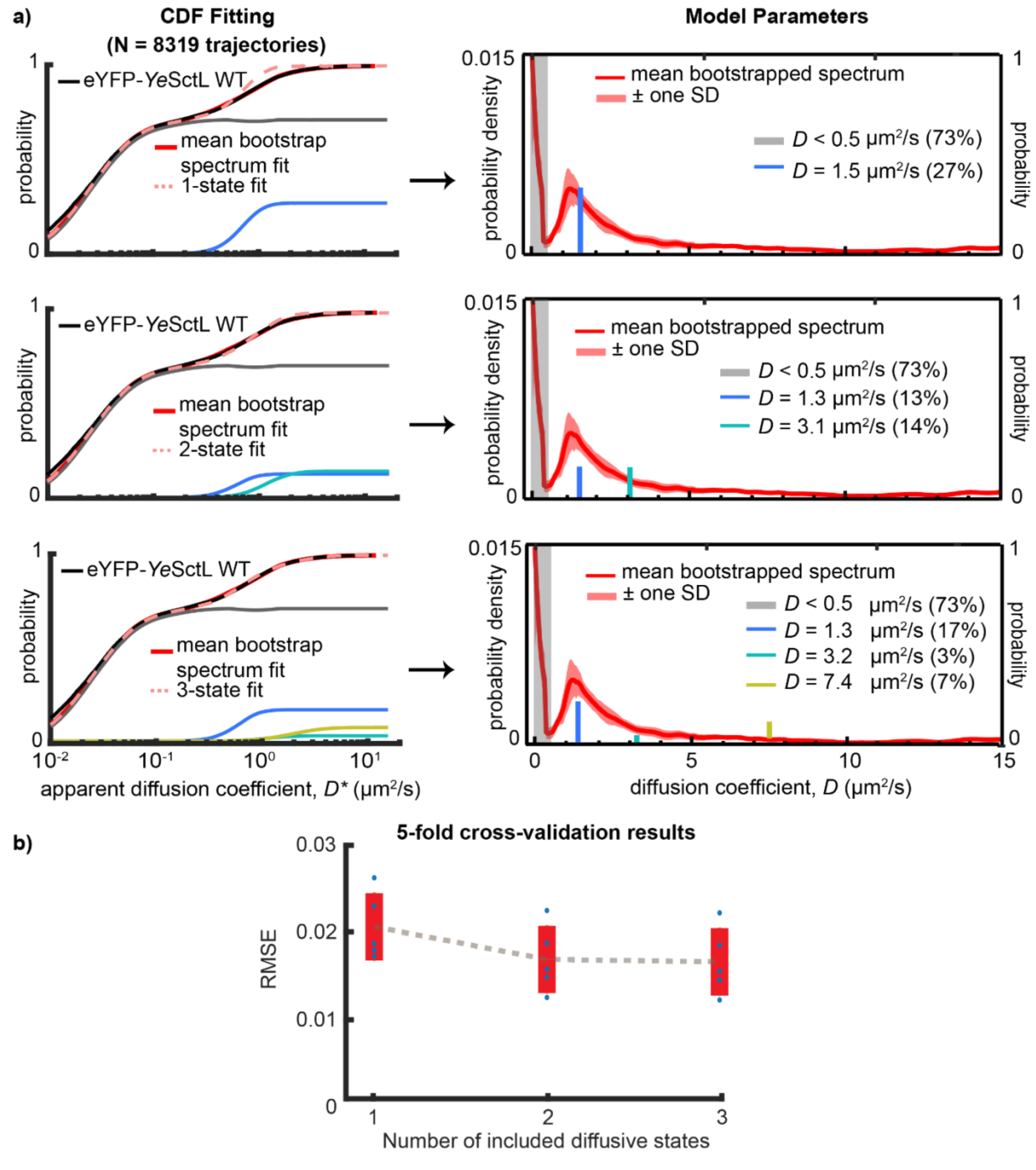


**Supplementary Figure 6:** a) Fitting model comparison for eYFP-*Ye*SctL single-molecule tracking data. Individual rows correspond to 1-, 2-, and 3-state models, respectively. A 2-state model converged on $D$ ~ 1.3 and 3.1μm$^2$/s, which agrees well with peaks in the diffusion coefficient spectrum. b) The 2-state model is also supported by 5-fold cross-validation analysis, which shows that a 2-state model does not overfit the data. A 3-state model could also have been chosen based on the cross-validation analysis. However, the 3-state model includes diffusive states at $D$ ~ 7.4 μm$^2$/s, which does not match the diffusion coefficient spectrum.

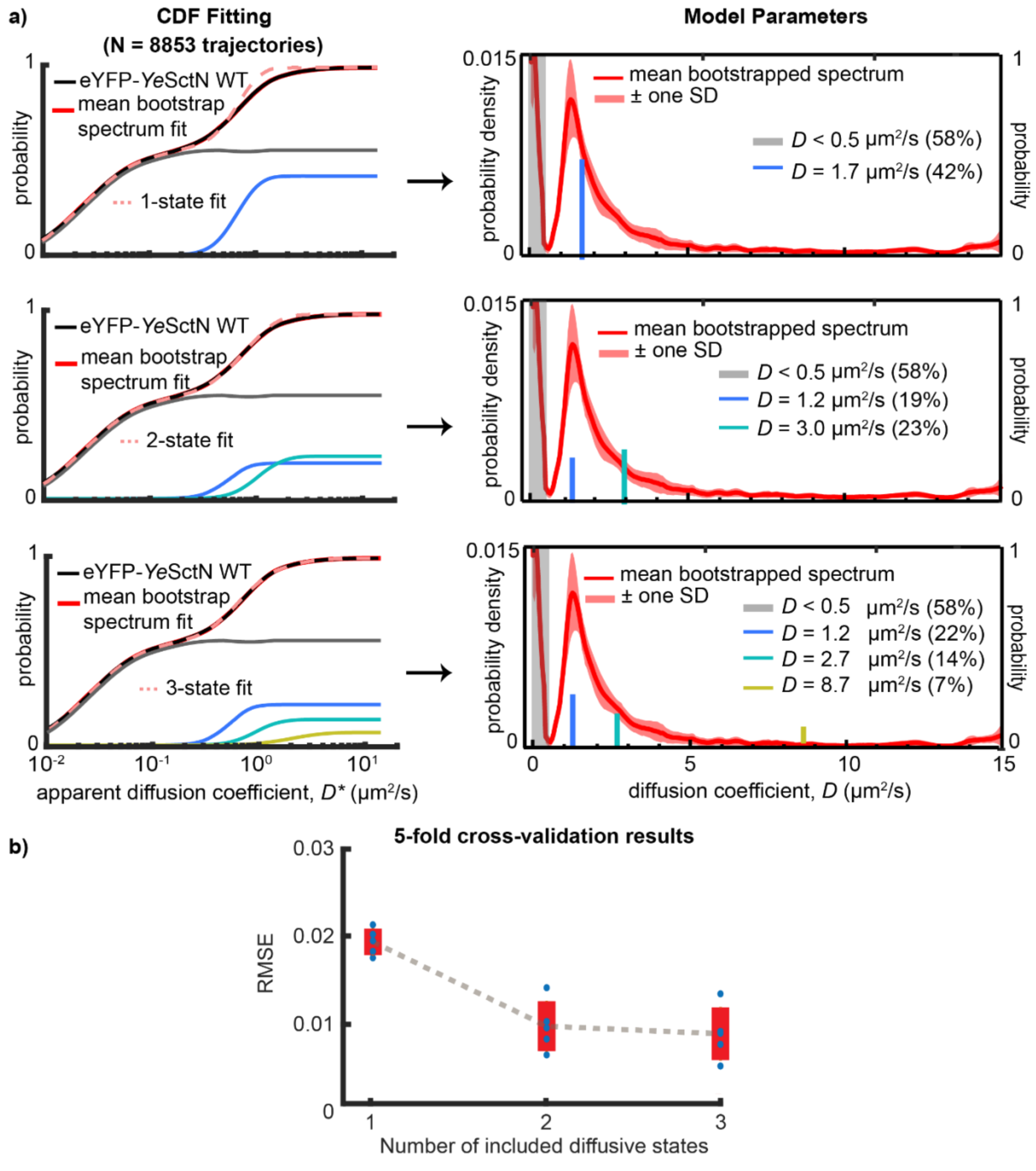


**Supplementary Figure 7:** a) Fitting model comparison for eYFP-*Ye*SctN single-molecule tracking data. Individual rows correspond to 1-, 2-, and 3-state models, respectively. A 2-state model converged on $D$ ~ 1.2 and 3.0 μm$^2$/s, which agrees well with peaks in the diffusion coefficient spectrum. b) The 2-state model is also supported by 5-fold cross-validation analysis, which shows that a 2-state model does not overfit the data. A 3-state model could also have been chosen based on the cross-validation analysis. However, the 3-state model includes diffusive states at $D$ ~ 8.7 μm$^2$/s, which does not match the diffusion coefficient spectrum.

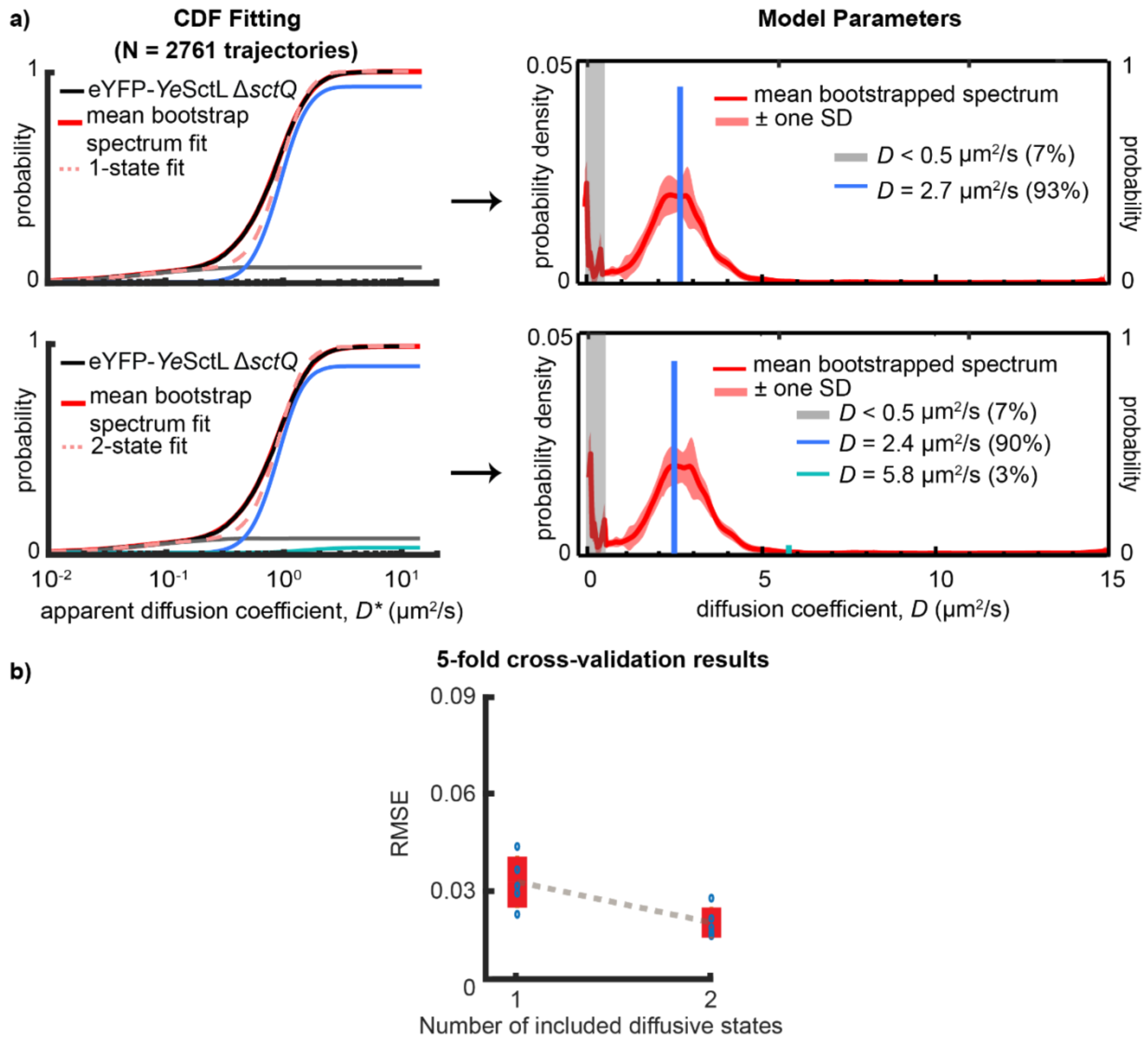


**Supplementary Figure 8:** a) Fitting model comparison for eYFP-*Ye*SctL Δ*sctQ* single-molecule tracking data. Individual rows correspond to 1-and 2-state models, respectively. A 1-state model converged on *D* ~ 2.7 μm$^2$/s, which agrees well with peaks in the diffusion coefficient spectrum. b) The 1-state model is also supported by 5-fold cross-validation analysis, which shows that a 1-state model does not overfit the data. A 2-state model could also have been chosen based on the cross-validation analysis. However, the 2-state model includes a diffusive state at *D* ~ 5.8 μm$^2$/s with a very small population fraction (3%) that does not agree with the diffusion coefficient spectrum. Thus, the 1-state model is favored over the 2-state model.

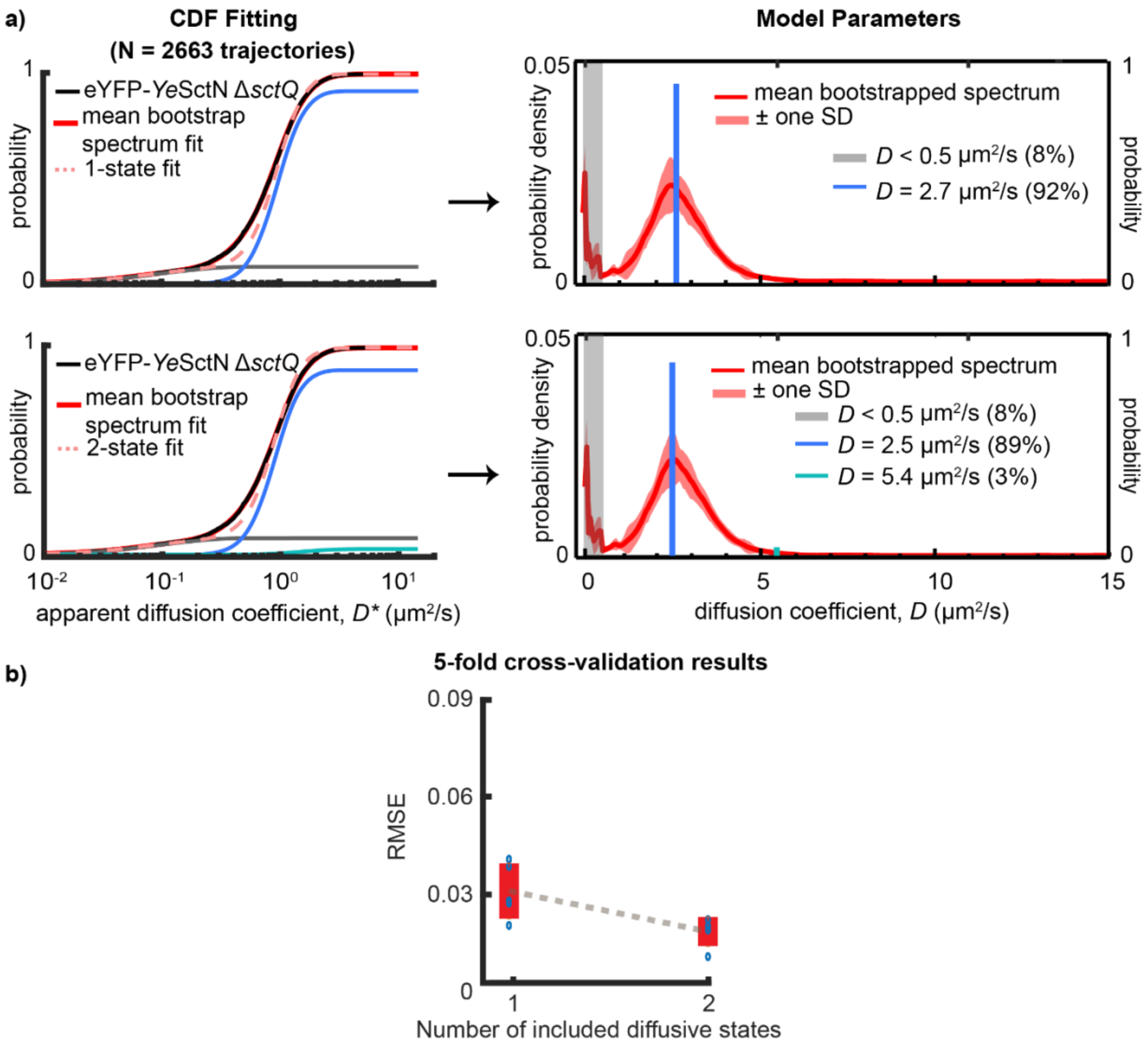


**Supplementary Figure 9:** a) Fitting model comparison for eYFP-*Ye*SctN Δ*sctQ* single-molecule tracking data. Individual rows correspond to 1-and 2-state models, respectively. A 1-state model converged on *D* ~ 2.7 μm$^2$/s, which agrees well with peaks in the diffusion coefficient spectrum. b) The 1-state model is also supported by 5-fold cross-validation analysis, which shows that a 1-state model does not overfit the data. A 2-state model could also have been chosen based on the cross-validation analysis. However, the 2-state model includes a diffusive state at *D* ~ 5.4 μm$^2$/s with a very small population fraction (3%) that does not agree with the diffusion coefficient spectrum. Thus, the 1-state model is favored over the 2-state model.

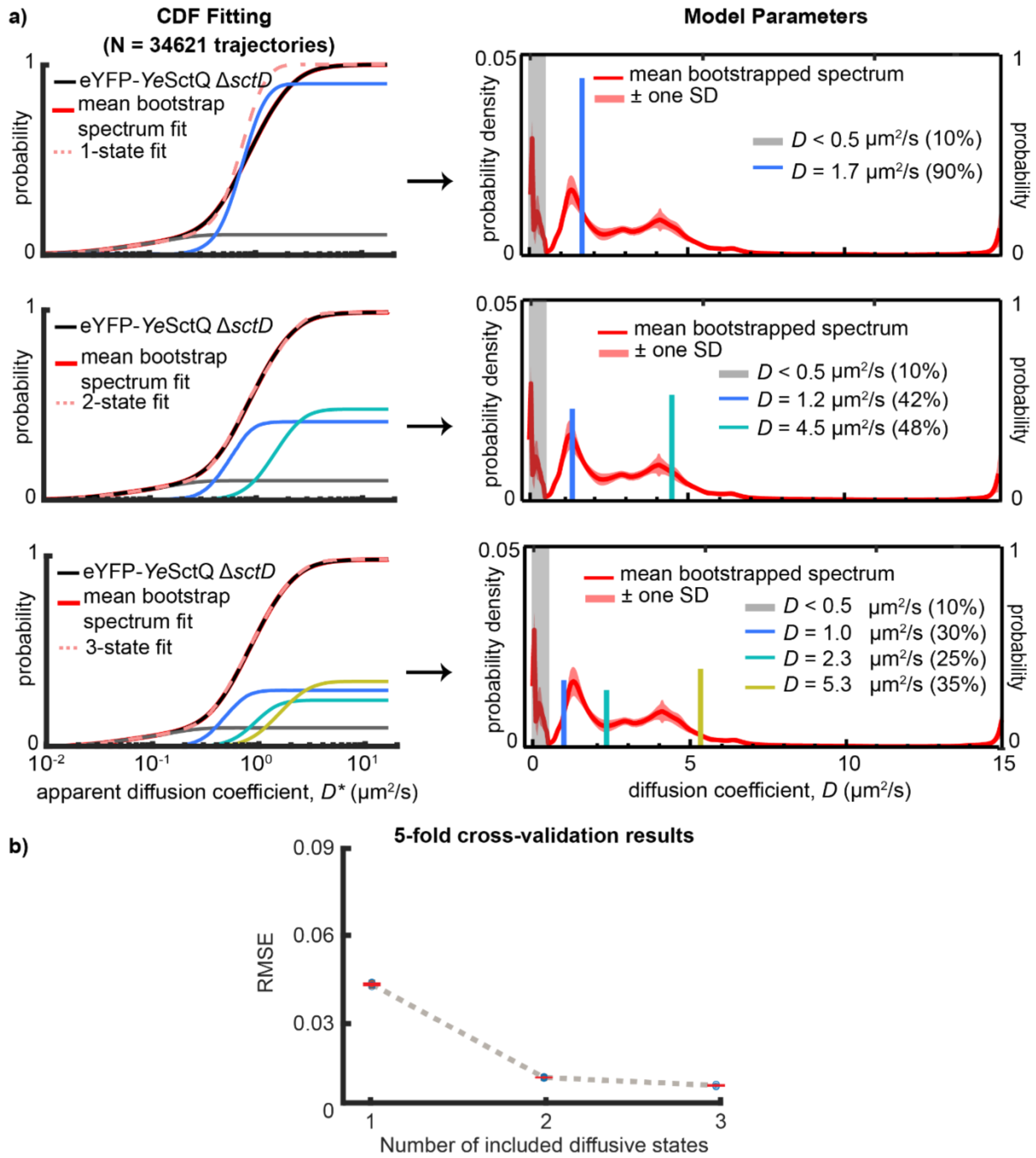


**Supplementary Figure 10:** a) Fitting model comparison for eYFP-*Ye*SctQ Δ*sctD* single-molecule tracking data. Individual rows correspond to 1-, 2- and 3-state models, respectively. A 2-state model converged on $D$ ~ 1.2 and 4.5 μm$^2$/s, which agrees well with peaks in the diffusion coefficient spectrum. b) The 2-state model is also supported by 5-fold cross-validation analysis, which shows that a 2-state model does not overfit the data. A 3-state model could also have been chosen based on the cross-validation analysis. However, the 3-state model includes a diffusive state at $D$ ~ 2.3 μm$^2$/s that does not agree with the diffusion coefficient spectrum. Thus, the 2-state model is favored over the 3-state model.

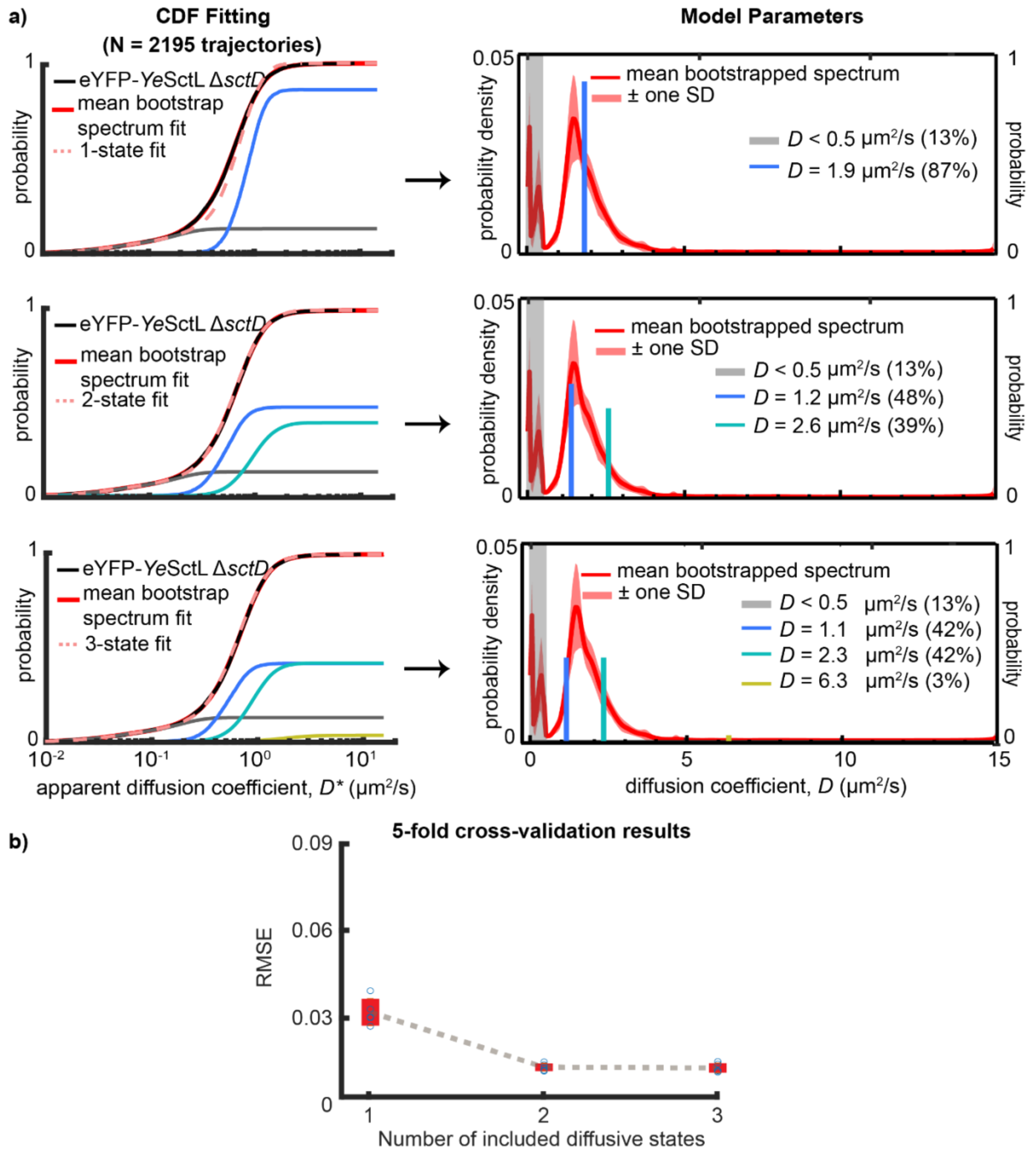


**Supplementary Figure 11:** a) Fitting model comparison for eYFP-*Ye*SctL Δ*sctD* single-molecule tracking data. Individual rows correspond to 1-, 2-, and 3-state models, respectively. A 2-state model converged on $D$ ~ 1.2 and 2.6 μm$^2$/s, which agrees well with peaks and spectral densities in the diffusion coefficient spectrum. b) The 2-state model is also supported by 5-fold cross-validation analysis, which shows that a 2-state model does not overfit the data. A 3-state model could have also been chosen based on the cross-validation analysis. However, the 3-state model includes a diffusive state at $D$ ~ 6.3 μm$^2$/s with a very small population fraction (3%) that does not agree with the diffusion coefficient spectrum. Thus, the 2-state model is favored over the 3-state model.

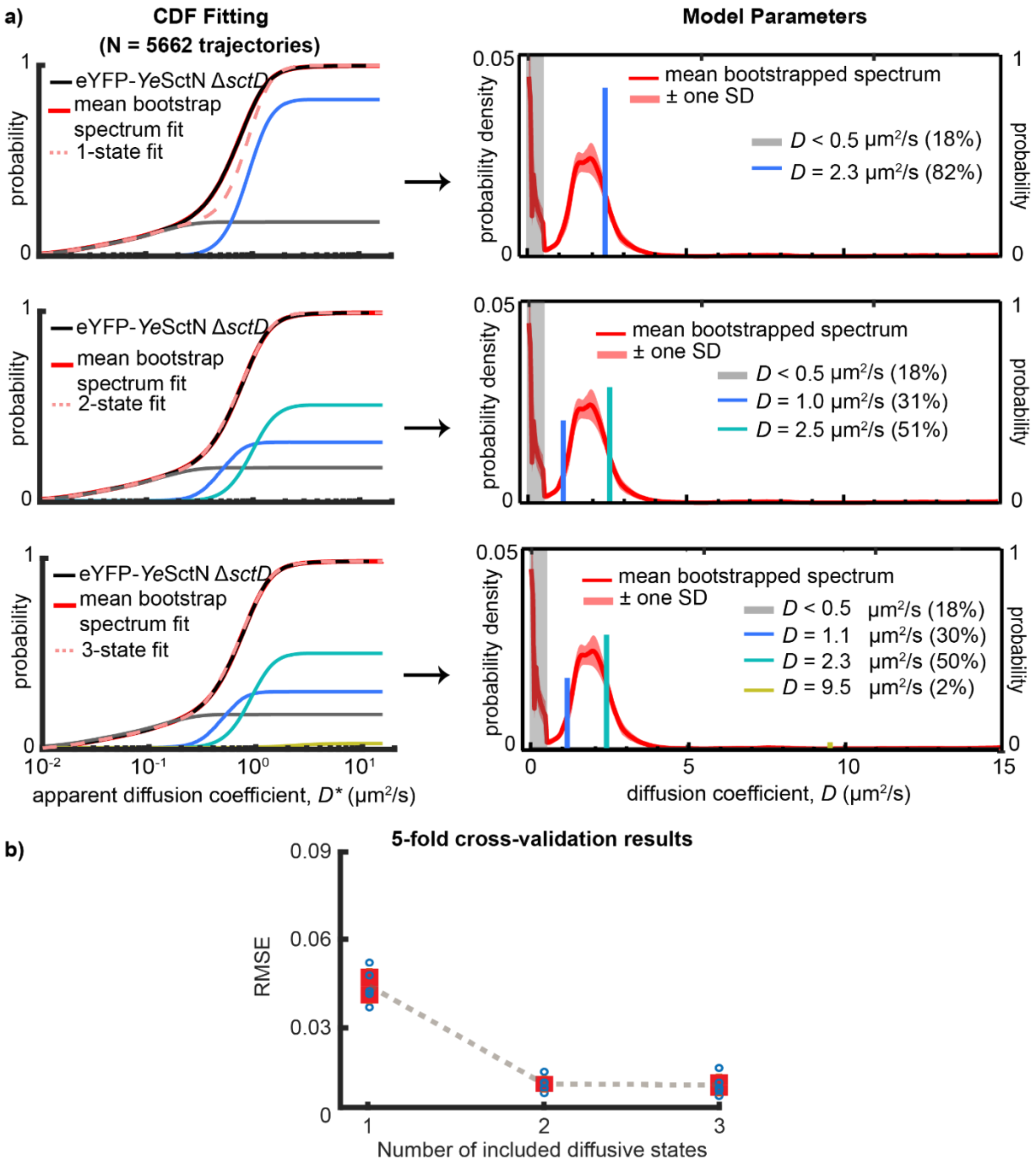


**Supplementary Figure 12:** a) Fitting model comparison for eYFP-*Ye*SctN Δ*sctD* single-molecule tracking data. Individual rows correspond to 1-, 2-, and 3-state models, respectively. A 2-state model converged on $D$ ~ 1.0 and 2.5 μm$^2$/s, which agrees well with peaks in the diffusion coefficient spectrum. b) The 2-state model is also supported by 5-fold cross-validation analysis, which shows that a 2-state model does not overfit the data. A 3-state model could have also been chosen based on the cross-validation analysis. However, the 3-state model includes a diffusive state at $D$ ~ 9.5 μm$^2$/s with a very small population fraction (2%) that does not agree with the diffusion coefficient spectrum. Thus, the 2-state model is favored over the 3-state model.

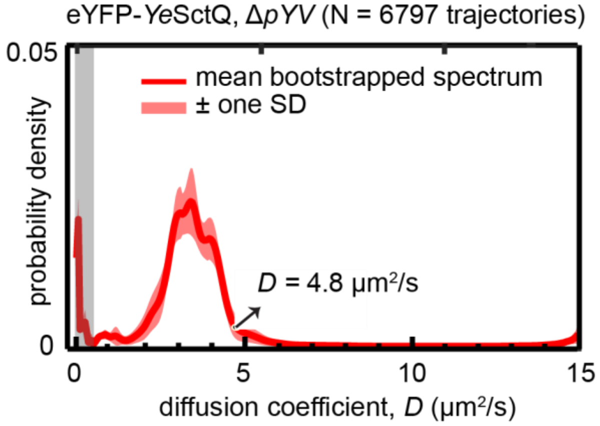


**Supplementary Fig. 13:** Diffusion coefficient spectrum for eYFP-*Ye*SctQ in the Δ*pYV* mutant showing spectral density between *D* ~ 3 to 5 μm$^2$/s. Δ*pYV* mutants do not express any other T3SS proteins.
